# A Comparison of Methods: Normalizing High-Throughput RNA Sequencing Data

**DOI:** 10.1101/026062

**Authors:** Rahul Reddy

## Abstract

As RNA-Seq and other high-throughput sequencing grow in use and remain critical for gene expression studies, technical variability in counts data impedes studies of differential expression studies, data across samples and experiments, or reproducing results. Studies like *Dillies et al. (2013)* compare several between-lane normalization methods involving scaling factors, while *Hansen et al. (2012)* and *Risso et al. (2014)* propose methods that correct for sample-specific bias or use sets of control genes to isolate and remove technical variability. This paper evaluates four normalization methods in terms of reducing *intra-group*, technical variability and facilitating differential expression analysis or other research where the biological, *inter-group* variability is of interest. To this end, the four methods were evaluated in differential expression analysis between data from *Pickrell et al. (2010)* and *Montgomery et al. (2010)* and between simulated data modeled on these two datasets. Though the between-lane scaling factor methods perform worse on real data sets, they are much stronger for simulated data. We cannot reject the recommendation of Dillies et al. to use TMM and DESeq normalization, but further study of power to detect effects of different size under each normalization method is merited.

## INTRODUCTION

RNA-Seq and other high-throughput sequencing technologies are key to the study of gene expression and especially important for differential expression analysis. However, technical variability in RNA-Seq data is an impediment to studying differential expression, reproducing results of experiments, and comparing raw count data between samples. For differential expression analysis in particular, our interest is in the biological variability between groups, rather than *intra-group,* technical variability that may arise during sequencing. Though comparative evaluations have been done using real data by *Hansen et al. (2012)*, and *Dillies et al. (2013)* have used both real and simulated data, it remains unclear which RNA-Seq normalization method(s) scientists should use for data analysis. More recently, *Risso et al. (2014)* proposed using factor analysis on sets of control genes, which also merits consideration. By comparing such recommended methods with real and simulated data, this paper seeks to help scientists choose the best normalization method for their own RNA-Seq analysis.

### Objective

This study will evaluate 4 normalization methods recommended for differential expression analysis in recent literature: Trimmed Mean of M-Values (TMM), Relative Log Expression (RLE, the default method in the DESeq package), Conditional Quantile Regression (CQN), and Remove Unwanted Variation (RUV). Using real and simulated data, these methods will be compared with respect to removing technical variability and differential expression analysis.

## METHODS

This section describes the normalization methods, real datasets, simulation, and criteria for comparison used in this study. In the following pages, terminology and abbreviations are used which are specific to Genomics, Statistics, the R statistical computing software, and the Bioconductor suite of R packages. For example, by technical variability we speak of variation *within* a group or condition, as opposed to the biological variation *between* groups or conditions that is of research interest. *Differential expression* refers to genes being up-regulated or down-regulated between different conditions. The condition may be that samples are from different individuals, populations, or species, different organ tissues or cell types for the same individuals, different time periods for the same individual and tissue, or any combination thereof. Typically, Bioconductor packages EdgeR or DESeq are the backbone of a differential expression study, as outlined by *Anders et al. (2013)*. This study uses the EdgeR package in a reference workflow provided by Susan Holmes (Stanford University, 2015), and implementing each method per its vignette.

### Normalization Methods

Most RNA-Seq datasets are obtained using Illumina sequencers that eight independent sequencing areas, or “lanes.” Libraries of cDNAs, representative of RNA from a cell or tissue sample, are deposited on these lanes. The normalization methods compared in this study address different sources of variation in the resulting count data, with the first two addressing differences in library size (i.e. sequencing depth). Library size, or number of mapped short reads obtained in sequencing a given library, can be normalized *between samples* by scaling raw read counts in each lane according to a single, lane-specific factor. Trimmed Mean of M-Values (TMM) and DESeq (Relative Log Expression, or RLE) are two different methods based on calculating such scaling factors. Conditional Quantile Normalization (CQN) corrects for within-sample, gene-specific effects (gene length, GC-content, etc.), while Remove Unwanted Variation (RUV) removes technical variability through factor analysis on sets of control genes or samples.

#### Trimmed Mean of M-Values (TMM)

TMM normalization is the EdgeR package’s default normalization method and assumes that most genes are not differentially expressed. It calculates a normalization factor for each gene, though this correction factor is applied to library size (i.e. sequencing depth) rather than not raw counts. Moreover, to compute the TMM factor, one lane is considered a reference sample and the others test samples, with TMM being the weighted mean of log ratios between test and reference, after excluding the most expressed genes and the genes with the largest log ratios. According to *Dillies et al. (2013)*, the DESeq method and TMM outperformed Median Count, Upper Quantile, Mean of Total Count, and RPKM (Reads per Kilobase per Million) normalization methods, as well as Quantile Normalization.

#### Relative Log Expression (DESeq)

For DESeq’s default normalization method, the RLE (Relative Log Expression) normalization method was used in EdgeR as it is equivalent. Starting with the hypothesis that most genes are not DE, scaling factors are calculated for each lane as median of the ratio, for each gene, of its read count of its geometric mean across all lanes. This way, non-differentially expressed genes will have similar read counts across samples, with a ratio of 1, and the median of the ratio for a given lane serves as a correction factor to apply to all read counts. Along with TMM, *Dillies et al. (2013)* recommended DESeq normalization.

#### Conditional Quantile Normalization (CQN)

Proposed by *Hansen et al. (2012)*, CQN operates on the assumption that RNA-Seq data is greatly affected by bias and systemic errors, with a sample effect that is not common for all genes. The authors focused on sample-specific GC-content bias, which resulted in confounding of GC-content and log fold change values. To remove such technical variability, they removed this GC-content effect and also used quantile normalization. However, they improved upon prior efforts to remove sample-specific GC bias by assuming that the distribution of true expression values does not depend on GC-content, and felt that strong GC-content effects across different parts of the same gene support this assumption. The authors also published a Bioconductor package, cqn, which we use to implement this normalization and create offsets that can be used in EdgeR’s GLM test (Likelihood Ratio Test) for differential expression.

#### Remove Unwanted Variation (RUV)

RUV is a normalization method that uses factor analysis of suitable sets of control genes (or samples) to remove technical variability, thus obtaining more accurate estimates of differential expression. *Risso et al. (2014)* used the External RNA Control Consortium’s spike-in controls as control genes and developed three algorithms. RUVg assumes a set of “negative” control genes that are not affected by biological covariates of interest (e.g. housekeeping genes or spike-ins), but are affected by unwanted technical variability like any other genes. RUVr makes no such assumption, and all control genes may be used to normalize the data while RUVs is a middle option that requires negative control genes exactly as in RUVg, but is less sensitive to these genes. Given housekeeping genes in our dataset that could serve as negative control genes, we implemented RUVg using the authors’ RUVSeq package.

### Real Data

#### Dataset Descriptions

Through the *tweeDESeqCountData* package by *Gonzalez (2010)* based on the study by *Esnaola et al. (2013),* we obtained the datasets of *Pickrell et al. (2010)* and *Montgomery et al. (2010)*. These studies of lymphoblastoid cell gene expression gathered RNA-Seq data from 69 individuals from Nigeria of Yoruba ancestry (*Pickrell*, HapMap YRI) and 60 from Utah of European ancestry (*Montgomery,* HapMap CEU). *Pickrell* included 29 men and 40 women, *Montgomery* 27 men and 33 women, and both datasets included GC-content and length data. No genes were known to be differentially expressed between populations, housekeeping genes were also included.

#### Comparison Procedures

The four methods were compared based on variation in normalized data results of differential expression analysis. After filtering out genes with low read counts, only 6855 genes were ultimately considered. Bar plots of the Biological Coefficient of Variance (BCV, square root of dispersion), violin plots of *intra-group* variation, and RLE (Relative Log Expression) Plots were made to see how each method reduced technical variability. Differential expression of genes between the two populations was determined using EdgeR’s GLM Likelihood Ratio Test, generating lists of genes obtained under each normalization method. These lists were then compared through a Venn Diagram and tables.

### Simulation

In addition to working with real data, we created a simulated dataset based on 6855 genes and 129 samples from the *Pickrell* and *Montgomery* data used in analysis. The SimSeq package allowed us to model RNA-Seq data, with read counts having a negative binomial distribution, generating count data for 6850 genes across 30 samples (15 per group) with 4000 of the genes differentially expressed. For this simulated dataset, we repeated the same analyses as with real data, and were also able to evaluate each method based on its false positive and false negative rates (or true positive and true negative rates). Having specified a priori that 4000 genes be randomly set as differentially expressed, we could measure each normalization method’s performance in finding these genes to be differentially expressed.

## RESULTS

Our interest in comparing normalization methods was to reduce technical, *intra-group* variability, and make biological, *inter-group* variability of research interest more apparent. For this, we considered real and simulated data. Looking at bar plots of the Biological Coefficient of Variation (BCV), TMM and DESeq hardly reduce the total variability (sum of intra- and inter-group) versus Raw Counts, though CQN and RUV seem to do so. We see this same pattern when considering only housekeeping genes, which have constant expression levels in most settings, though we cannot make any statements about significance.

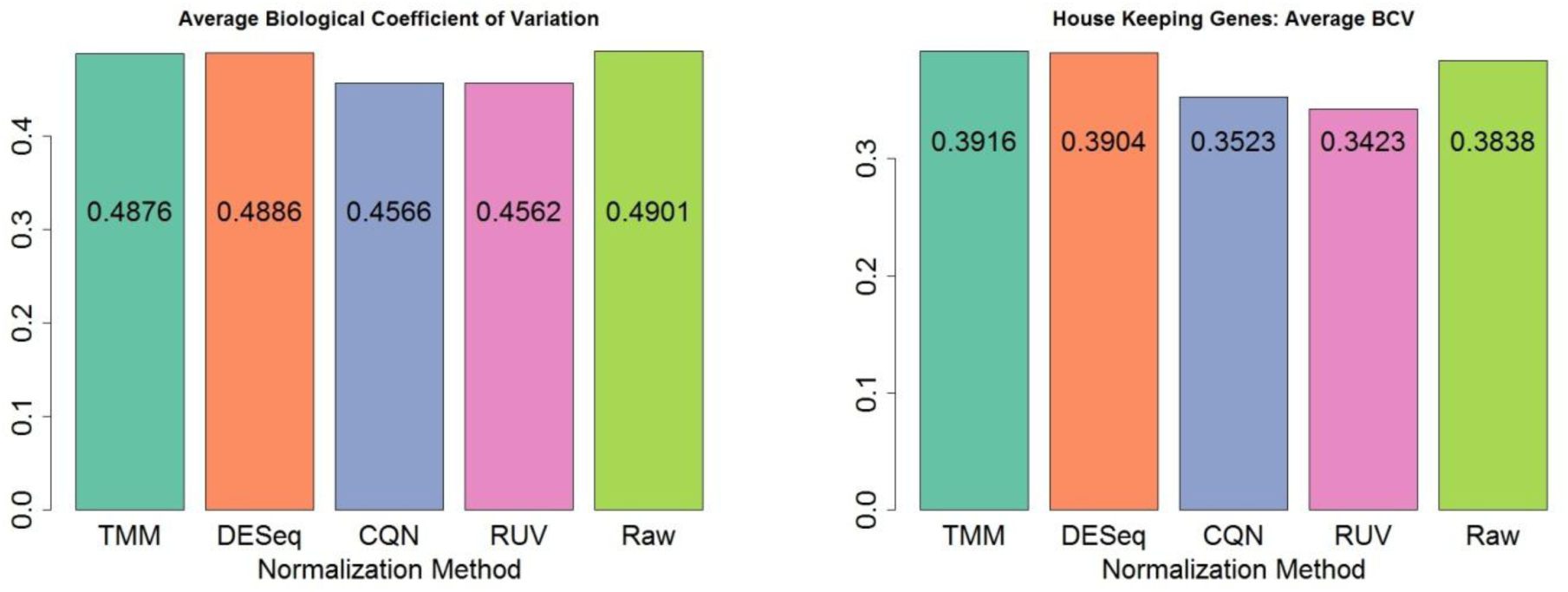

Though including CQN and RUV was not possible in the violin plots below, we see that TMM and DESeq again reduce variation (within each group, i.e. population) to only a limited degree.

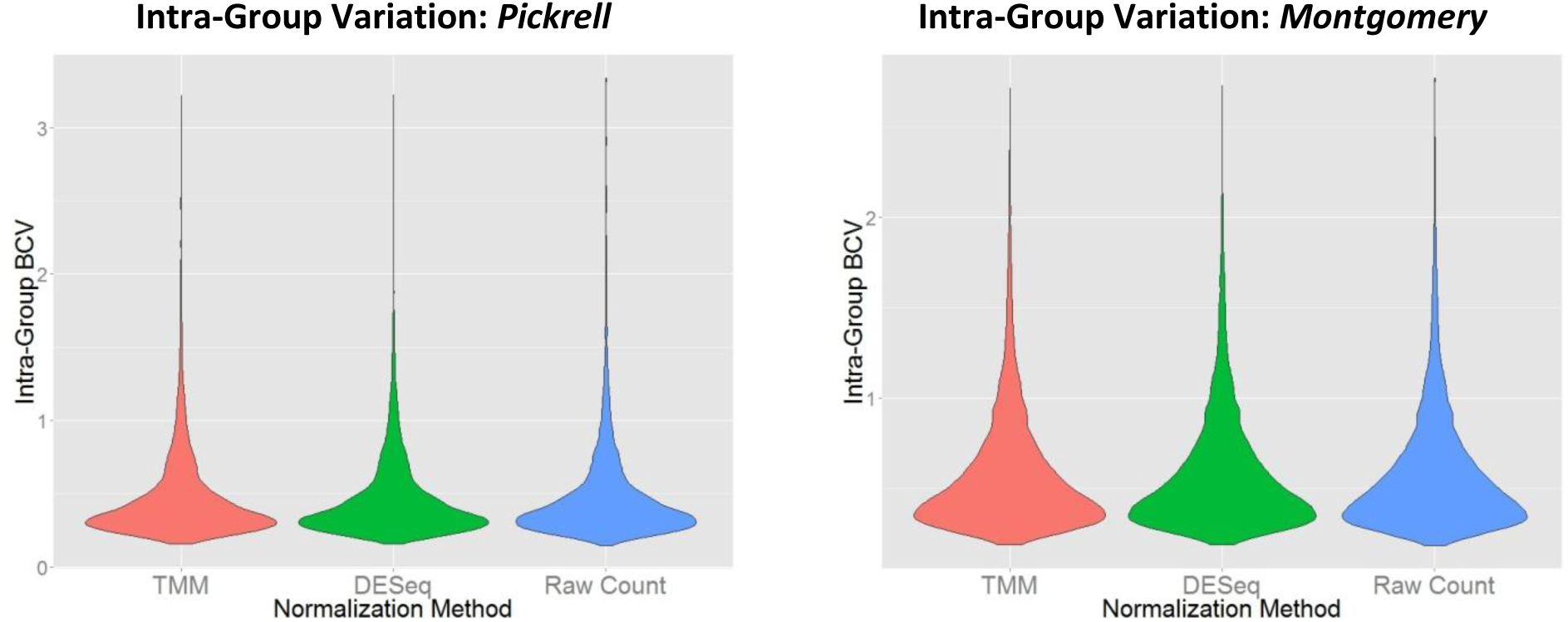

Looking at Relative Log Expression (RLE) plots for the full dataset, RUV clearly reduces much of the *intra-group* variation. Its use of housekeeping genes as negative controls may explain reduced *intra-group* variation in particular, as housekeeping genes should not change expression levels across conditions.

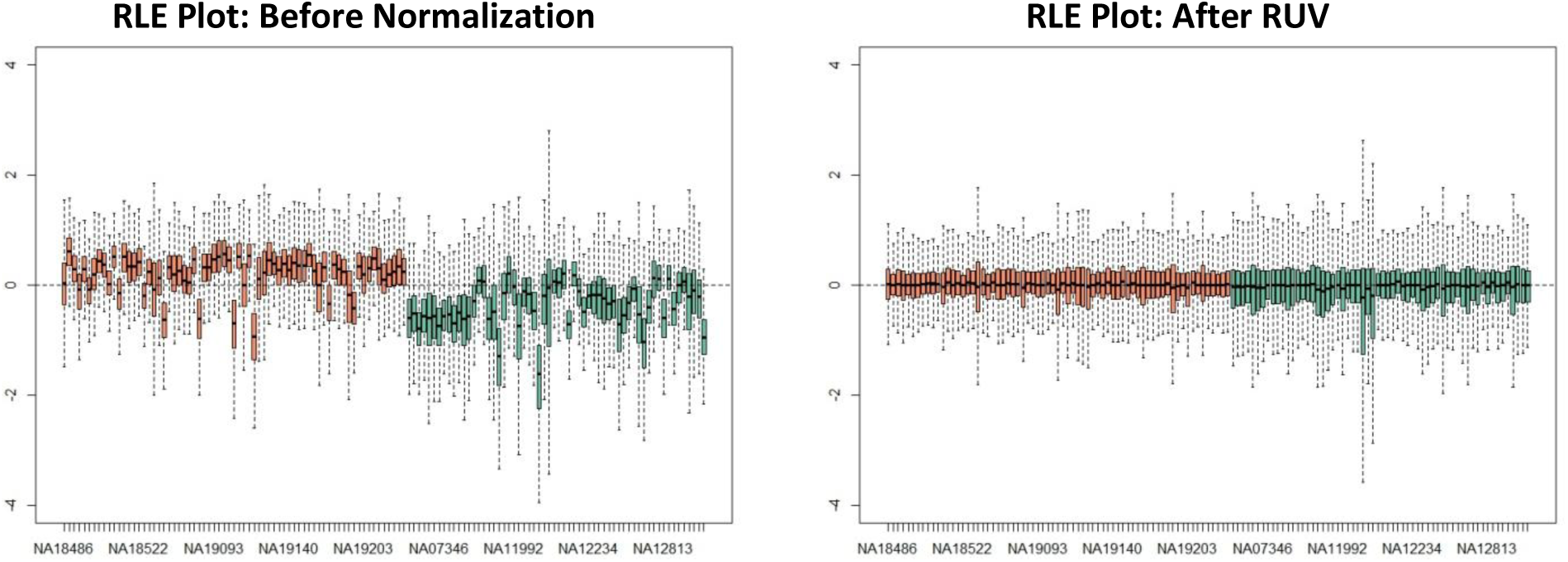
(*Pickrell’s* 69 samples are on the left side of the RLE plots in red, *Montgomery’s* 60 samples are on right in green)

Furthermore, in differential expression results, RUV and CQN yielded the largest counts (4384 and 4371 respectively), while TMM and DESeq yielded fewer (4207 and 4194 respectively), with 3240 found under all four methods. CQN also led to 911 genes listed as differentially expressed that were not found under any other method. The following table and Venn Diagram show counts of differentially expressed genes and those found in common for each method.

(6855 genes analyzed for differential expression, after filtering out the 45725 genes that did not have at least 3 or more samples with at least 10 reads

Differentially Expressed Genes by Method
|  | TMM | DESeq | CQN | RUV | Raw |
| --- | --- | --- | --- | --- | --- |
| TMM | 4207 | 4075 | 3339 | 4165 | 3872 |
| DESeq | - | 4194 | 3292 | 4087 | 3989 |
| CQN | - | - | 4371 | 3410 | 3192 |
| RUV | - | - | - | 4384 | 3911 |
| Raw | - | - | - | - | 4154 |

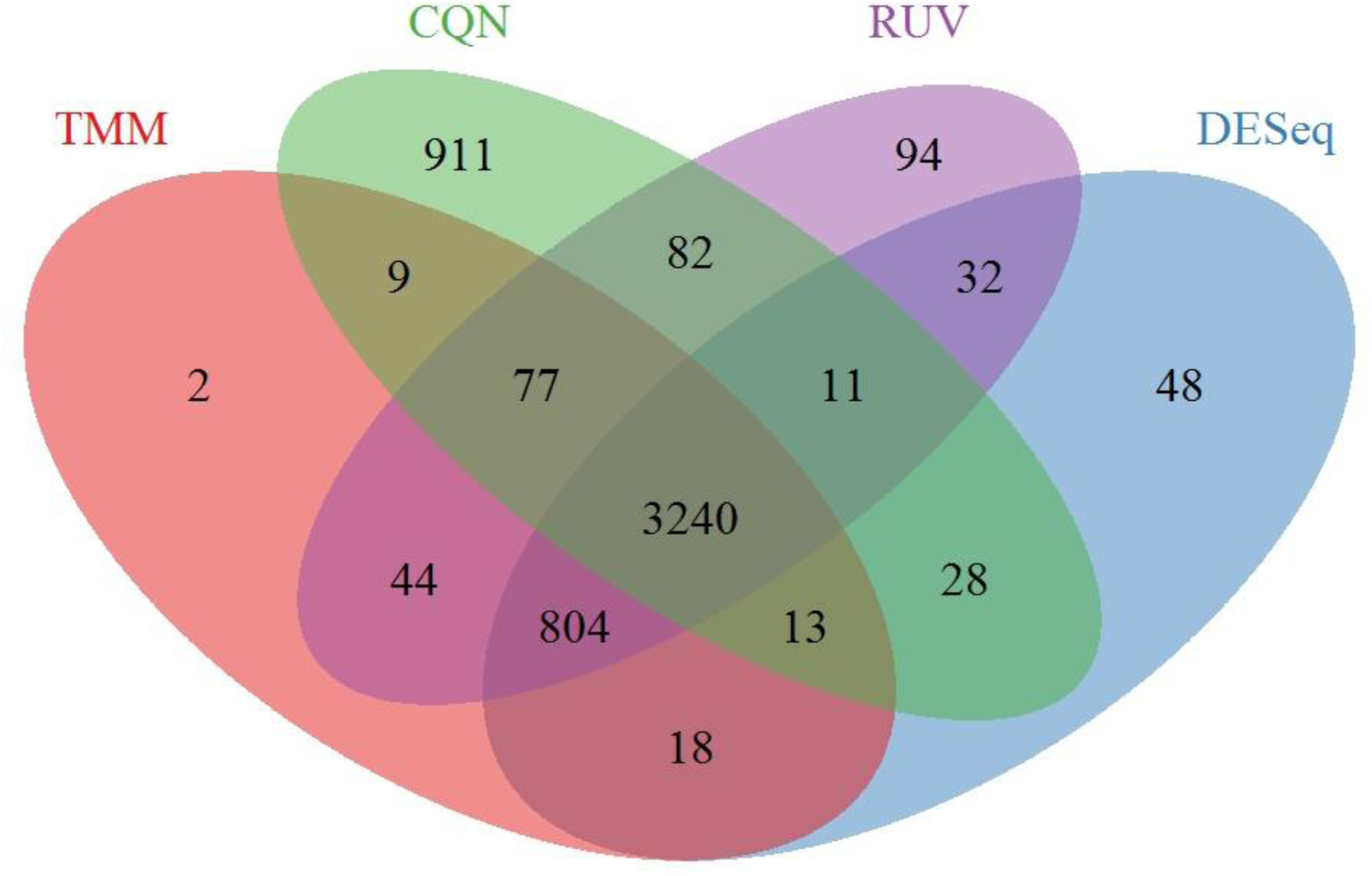
**Venn Diagram: Differentially expressed genes found in common for each normalization method**

Making further claims is difficult though, as no genes are known to be differentially expressed across our two populations, in the manner of housekeeping genes’ constant expression across cells and conditions.

### Simulation

Simulation helped assess out-of-sample effect on technical, *intra-group* variability and measure error rates in detecting differential expression for each method. Variability is again barely affected by our normalization methods, save for RUV:

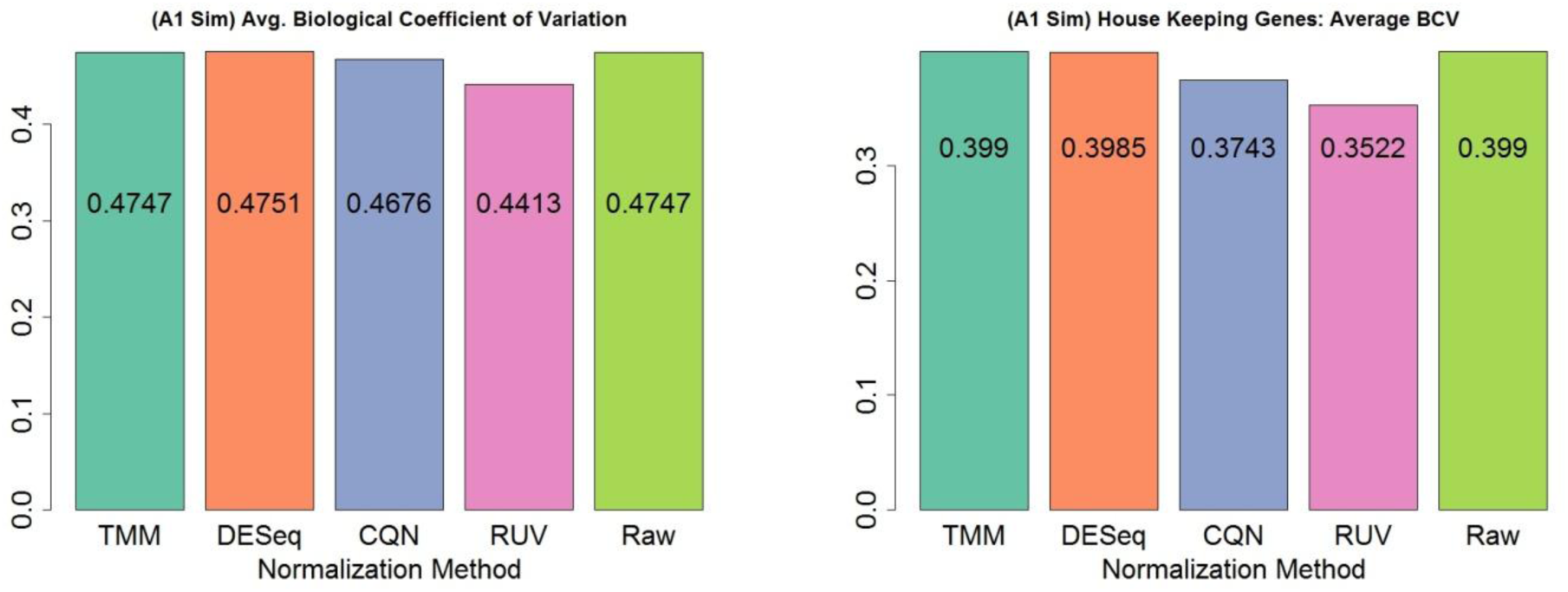

Considering the RLE plots for simulated data, RUV again visibly reduces intra-group variance, though this is a smaller sample size (30 individuals, 15 per group) due to the nature of the SimSeq package.

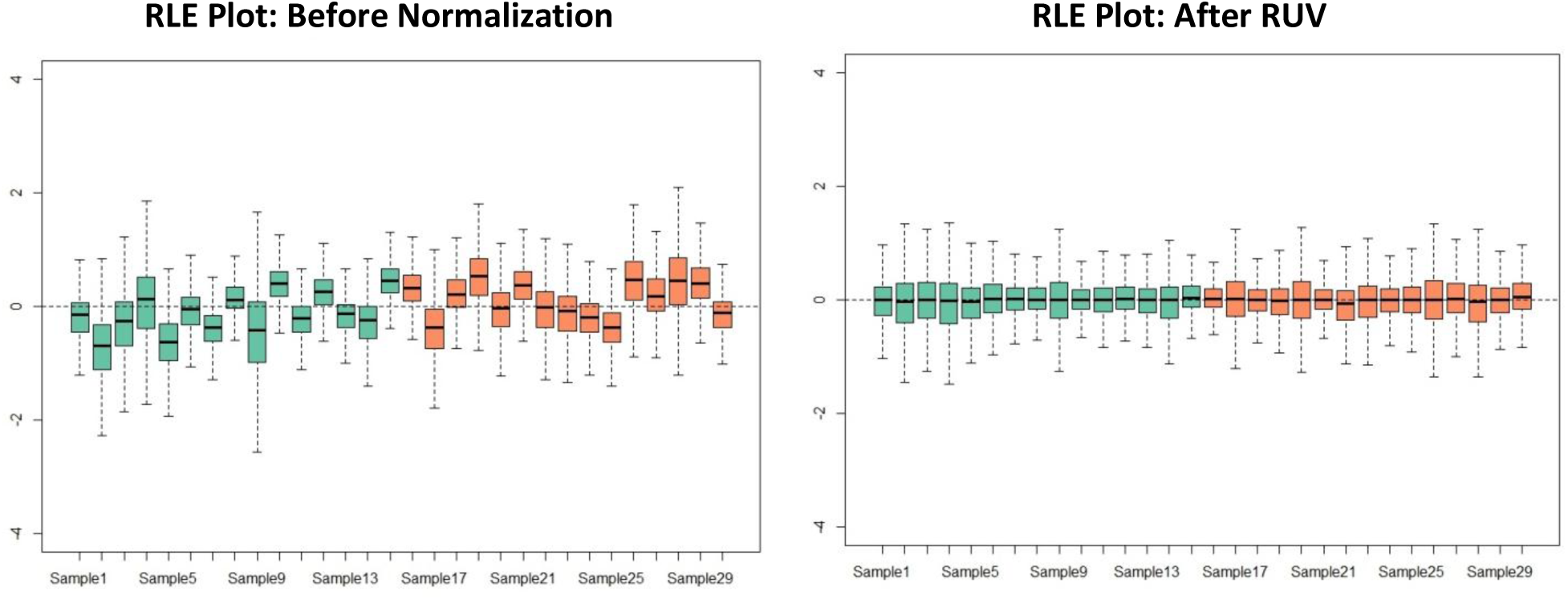

Differentially Expressed Genes by Method
|  | TMM | DESe | CQN | RUV | Raw |
| --- | --- | --- | --- | --- | --- |
| **TMM** | 1327 | 1318 | 870 | 726 | 1327 |
| **DESe** | - | 1337 | 874 | 723 | 1318 |
| **CQN** | - | - | 1098 | 647 | 870 |
| **RUV** | - | - | - | 864 | 726 |
| **Raw** | - | - | - | - | 1327 |

Of 6850 simulated genes, 4000 were set at random to be differentially expressed. After filtering, 6183 genes remained for our analysis. The Venn Diagram on the right gives a breakdown of differential expression by method, while the *accuracy* of each normalization method is given in the following confusion matrices. The error rate is given for each method, with (in clockwise order from top left) True Positive, False Negative, True Negative, and False Positive counts.

| TM | 41%<br>Error | DE Analysis |  |
| --- | --- | --- | --- |
|  |  | DE | Not DE |
| Real | DE | 1219 | 2427 |
|  | Not DE | 108 | 2429 |

Differential expression results were much smaller in counts, but also notable for CQN and RUV having the smallest counts, whereas they had the most for the real data. simulation also allowed us to compare the four methods by their false positive rate in differential expression analysis.

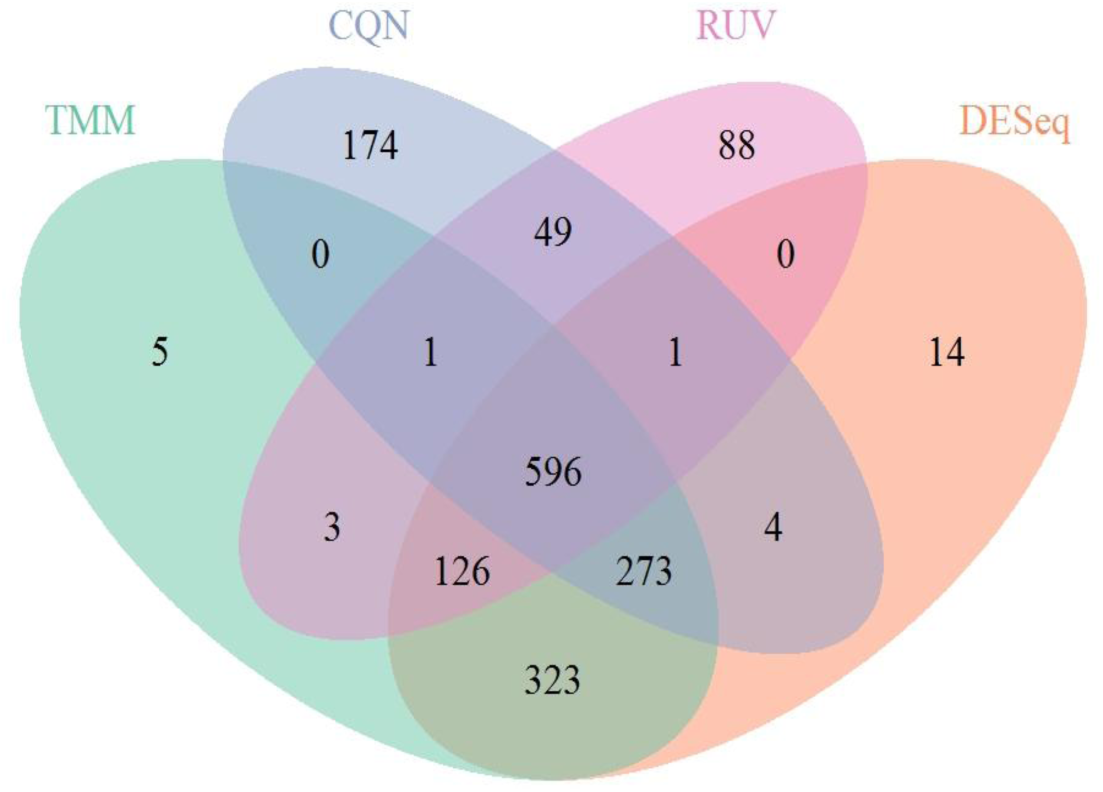
**DE genes found in common for each method**

| DESeq | 41%<br>Error | DE Analysis |  |
| --- | --- | --- | --- |
|  |  | DE | Not DE |
| Real | DE | 1224 | 2422 |
|  | Not DE | 113 | 2424 |

|  | CQN | 43.09%<br>Error | DE Analysis |  |
| --- | --- | --- | --- | --- |
|  |  |  | DE | Not DE |
| Real | DE |  | 1040 | 2606 |
|  | Not DE |  | 58 | 2479 |

|  | RUV | 46.06%<br>Error | DE Analysis |  |
| --- | --- | --- | --- | --- |
|  |  |  | DE | Not DE |
| Real | DE |  | 831 | 2815 |
|  | Not DE |  | 33 | 2504 |

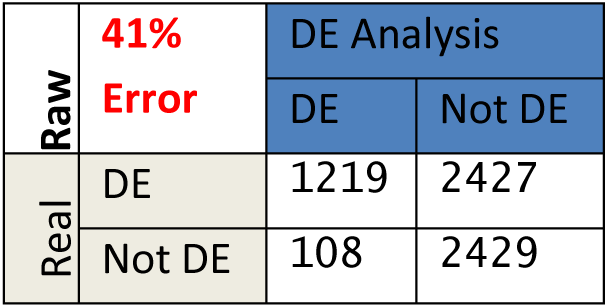

|  | Raw | 41%<br>Error | DE Analysis |  |
| --- | --- | --- | --- | --- |
|  |  |  | DE | Not DE |
| Real | DE |  | 1219 | 2427 |
|  | Not DE |  | 108 | 2429 |

From the simulated data, it’s clear that even though RUV seemed best in reducing intra-group variance with real data, it has a high false negative rate (2805 genes classified as Not DE, when in fact they were DE) and did worst of all the methods in differential expression analysis.

## DISCUSSION

In considering four methods to reduce technical, intra-group variability and thus improve differential expression analysis, there was not one method that dominated. In reducing intra-group, technical variability TMM and DESeq had no significant effect as compared to the Raw Count data, whereas CQN and RUV may have had an effect. However, CQN and RUV are only useful in particular circumstances. CQN can correct for GC-content, read lengths, or similar sample-specific bias, which is why *Dillies et al. (2013)* did not evaluate it against TMM and DESeq, the methods they found to perform best. RUV requires a set of negative control genes or samples, such as housekeeping genes, which are not always known. For the *Pickrell* and *Montgomery* datasets, CQN and RUV reduced variation more and resulted in a higher count of differentially expressed genes as well.

However, simulation suggests that these advantages of CQN and RUV may have been specific to our choice of datasets, and that much more can be done to assess how each normalization method affects the power of tests for differential expression. The *Pickrell* and *Montgomery* datasets would be expected to have non-trivial GC-content, as they were used by *Hansen et al. (2012)* to introduce CQN. They also had housekeeping genes, ready to use as controls for RUV. For differential expression as well, simulation results were the opposite of those with real data, as RUV and CQN had the lowest gene counts. Moreover, their error rates (46.09% and 43.03%) were higher as well, with very high false negative rates. Though our scope did not include the effect of each normalization method on power (i.e. minimizing false negatives), RUV’s high rate of false negatives and the ability with simulation to test power by varying the number of differential expressed genes make this worth future study.

### Conclusion

Though this study found CQN and RUV promising in their ability to reduce technical, *intra-group* variability, it cannot reject the recommendation of *Dillies et al. (2013)* to use DESeq and TMM normalization for differential expression analysis. Indeed, these two methods were the most accurate for simulated data and had similar differential expression results to CQN and RUV using real data. What remains of interest are the conditions under which CQN and RUV may be best applied, especially if certain levels of GC-content bias meriting use of CQN or specific control genes best suited to RUV could be identified. In the manner of a 2013 protocol established by Anders et al for RNA sequencing data analysis, protocols for normalization are worth developing as more and more research and researchers rely on RNA-Seq.

